# CRISPR-Cas13d induces efficient mRNA knock-down in animal embryos

**DOI:** 10.1101/2020.01.13.904763

**Authors:** Gopal Kushawah, Joaquin Abugattas-Nuñez del Prado, Juan R. Martinez-Morales, Michelle DeVore, Javier R. Guelfo, Emry O. Brannan, Wei Wang, Timothy J. Corbin, Andrea M. Moran, Alejandro Sánchez Alvarado, Edward Málaga-Trillo, Carter M. Takacs, Ariel A. Bazzini, Miguel A. Moreno-Mateos

**Affiliations:** Stowers Institute for Medical Research, 1000 E 50th St, Kansas City, MO 64110, USA; Andalusian Center for Developmental Biology (CABD), Pablo de Olavide University/CSIC/Junta de Andalucía, Ctra. Utrera Km.1, 41013, Seville, Spain; Department of Molecular Biology and Biochemical Engineer, Pablo de Olavide University, Ctra. Utrera Km.1, 41013, Seville, Spain; Department of Biology, Universidad Peruana Cayetano Heredia, Av. Honorio Delgado 430, Lima 15102, Perú; University of New Haven, West Haven, Connecticut 06516, USA; Howard Hughes Medical Institute, Stowers Institute for Medical Research, Kansas, MO, USA; Department of Molecular and Integrative Physiology, University of Kansas Medical Center, 3901 Rainbow Blvd, Kansas City, KS 66160, USA

**Keywords:** CRISPR-Cas13, embryogenesis, zebrafish, RNA targeting

## Abstract

Early embryonic development is driven exclusively by maternal gene products deposited into the oocyte. Although critical in establishing early developmental programs, maternal gene functions have remained elusive due to a paucity of techniques for their systematic disruption and assessment. CRISPR-Cas13 systems have recently been employed to induce RNA degradation in yeast, plants and mammalian cell lines. However, no systematic study of the potential of Cas13 has been carried out in an animal system. Here, we show that CRISPR-Cas13d is an effective and precise system to deplete specific mRNA transcripts in zebrafish embryos. We demonstrate that both zygotically-expressed and maternally-provided transcripts are efficiently targeted, resulting in an 80% average decrease in transcript level and the recapitulation of well-known embryonic phenotypes. Moreover, we show that this system can be used in medaka, killifish and mouse embryos. Altogether our results demonstrate that CRISPR-Cas13d is an efficient knock-down platform to interrogate gene function in animal embryos.

## Introduction

The experimental dissection of gene function has been radically transformed with the advent of DNA engineering technologies such as TALEN and CRISPR-Cas9 systems (Hsu et al., 2014). These tools allow researchers to link genotype to cellular phenotypes through the generation of permanent changes to the genome. Complementary ‘knock-down’ approaches such as RNA interference have also proven to be invaluable tools to interrogate gene function. However, in aquatic vertebrate organisms such as zebrafish (*Danio rerio*), other teleost fish and *Xenopus* (Chen et al., 2017; Lund et al., 2011), attempts to establish efficient RNAi technologies have largely failed (Chen et al., 2017; Kelly and Hurlstone, 2011; Lund et al., 2011). In their place, morpholinos (MOs), and more recently, antisense oligonucleotides (ASOs), have been used to perturb RNA activity (Pauli et al., 2015; Stainier et al., 2017). While an ASO-based approach has not been followed up since its implementation in zebrafish (Pauli et al., 2015), MOs have been widely implemented for over two decades. MOs are nucleic acid-analog antisense oligomers that disrupt RNA function/output by blocking translation or splicing (Stainier et al., 2017). However, the utility of MOs has recently been called into question due to observed toxicity and off-target effects (Gentsch et al., 2018; Joris et al., 2017; Kok et al., 2015; Lai et al., 2019; Robu et al., 2007; Schulte-Merker and Stainier, 2014). In several instances, discordant phenotypes have been observed between MO-treated animals and loss-of-function mutant animals and, more strikingly, additional phenotypes have been observed upon MO treatment in loss-of-function mutant backgrounds, indicating cellular effects unrelated to targeted RNA knock-down (Joris et al., 2017; Kok et al., 2015). More recently, studies have shown that some MOs can trigger an innate immunity response, interferon-stimulated and cellular stress response gene up-regulation, and off-targeting through mis-splicing in zebrafish and Xenopus (Gentsch et al., 2018; Joris et al., 2017; Lai et al., 2019; Robu et al., 2007), reinforcing the need for established controls to ensure the fidelity of MO-induced phenotypes (Stainier et al., 2017). These concerns, as well as the high expense of morpholinos, tedious methods for validating efficacy and fidelity (i.e. assessment of protein output or ribosome occupancy) (Lee et al., 2013), and the inability to implement in a tissue-specific or temporal manner, have limited their use in systematic approaches to studying gene function in teleost and amphibian embryos.

RNA ‘knock-down’ approaches hold many advantages compared to DNA mutagenesis (e.g. Cas9). The ability to directly disrupt gene activity sidesteps the need for laborious multi-generation genotype screening to establish permanent genetic strains. Likewise, knock-down strategies allow for the study of *in vivo* maternal effects without the need for homozygous mutant mothers (Ciruna et al., 2002; Moreno-Mateos et al., 2015), which for many critically important genes are either unviable or infertile (Abrams and Mullins, 2009; Ciruna et al., 2002). Furthermore, by directly manipulating RNA activity in a relatively tunable manner, researchers can study how subtle changes in transcript levels impact biological processes. In fact, by reducing but not completely removing gene activity, researchers can uncover phenotypes that may otherwise remain hidden due to loss-of-function lethality (Smith et al., 2017). Finally, therapeutic approaches that target RNA rather than DNA can be advantageous in providing temporary but effective modifications without the prospect of permanent heritable changes (Cox et al., 2017).

Cas13 is a class 2/type VI CRISPR-Cas RNA endonuclease, which has recently been employed to induce both the cleavage and subsequent degradation of RNA in fission yeast, plants and mammalian cell lines (Abudayyeh et al., 2017; Aman et al., 2018; Cox et al., 2017; Jing et al., 2018; Konermann et al., 2018). This method has been shown to be more effective and specific than RNAi in mammalian cells (Abudayyeh et al., 2017; Cox et al., 2017; Konermann et al., 2018). However, to date, no systematic study of the potential of Cas13 has been carried out in an animal model system. Here, we show that CRISPR-RfxCas13d (CasRx), but not Psp/PguCas13b or LwaCas13a, is an effective and precise system to deplete specific mRNA transcripts in zebrafish embryos. We demonstrate that zygotically-expressed, as well as ectopic or maternally-provided transcripts are efficiently targeted, resulting in an average of more than 80% decrease in transcript levels and the recapitulation of well-known maternal and/or zygotic embryonic phenotypes. Importantly, through whole transcriptome sequencing, we observed highly specific knock-down. Finally, we successfully implement CRISPR-Cas13d system in other research organisms such as medaka (*Oryzias latipes*), killifish (*Nothobranchius furzeri*) and mouse (*Mus musculus*) embryos, demonstrating its applicability across a range of aquatic and terrestrial animal models. Our results establish CRISPR-Cas13d as an efficient and straight-forward method for the systematic, tractable, and unambiguous study of gene function *in vivo* during embryogenesis across a range of animal species.

### Design

Gene knock-down approaches such as RNA interference serve an invaluable role in elucidating gene function. However, in aquatic vertebrate organisms such as zebrafish and other teleost fish, RNA interference has failed to develop as an efficient and systematic tool. Instead, MOs have been used to disrupt gene function by blocking translation or preventing RNA splicing. However, MOs can induce nonspecific effects that are separate from target RNA perturbation(Gentsch et al., 2018; Joris et al., 2017; Lai et al., 2019; Robu et al., 2007). Further, researchers have found that MO-based phenotypes can differ form those obtained by genetic mutation (Kok et al., 2015). Although these differences may be explained, in part, by genetic compensation (El-Brolosy et al., 2019; Ma et al., 2019; Rossi et al., 2015), the use of MOs requires several proper controls to ensure accuracy (Stainier et al., 2017). Based on the limitations and caveats of current knockdown technologies in teleosts, we set out to develop and optimize a Cas13/CRSPR-based method to disrupt gene function. We had two goals: to develop 1) a technique that could be used in a cost-efficient and systematic manner and 2) a viable approach to interrogate maternally-provided developmental programs in the early embryo. The study of early development in animal embryos, which is driven by thousands of mRNAs that are maternally provided (Lee et al., 2014), requires elaborate genetic schemes to remove maternal activity. Herein, we find that, among various Cas13 variants tested, RfxCas13d efficiently disrupts maternal and zygotic gene function without embryonic toxicity, and with high specificity, in zebrafish embryos. We successfully apply this approach to gene knockdown in medaka, killifish and mouse embryos. Therefore, CRISPR-RfxCas13d is a robust and cost-effective technology to systematically disrupt gene function in vertebrates. We envision the application of this method to the study of gene expression in a high-throughput manner in the early embryo, as well as applied for use in tissue-specific manners.

## Results

### Optimization of RNA targeting by CRISPR-Cas13 in zebrafish embryos

To assess the potential toxic effects of different Cas13 proteins successfully used in mammalian cells (Abudayyeh et al., 2017; Cox et al., 2017; Konermann et al., 2018), we constructed expression vectors to generate mRNA encoding four different Cas13 variants (*RfxCas13d, PguCas13b, PspCas13b* or *LwaCas13a-GFP*). The mRNA for each Cas13 variant was injected individually into one-cell stage zebrafish embryos (Figure 1A). Protein expression *in vivo* was confirmed for all Cas13 variants tested (Figure S1A) except LwaCas13a-GFP likely due to a low stability of *LwaCas13* mRNA as reported in mammalian cells (Abudayyeh et al., 2017) (Figure S1B). While PguCas13b and PspCas13b negatively impacted embryonic development (Figures 1B and S1C), RfxCas13d displayed no toxic effects (up to 300 pg of mRNA per embryo; Figure S1D). Hence, we limited subsequent analyses of endogenous mRNA abrogation to RfxCas13d. First, to test whether Rfx-Cas13d could trigger the degradation of specific mRNAs in zebrafish embryos, we designed six guide RNAs (gRNAs) complementary to different sequences within the coding sequence (CDS) of the *tbxta* mRNA pooled into 2 sets of gRNAs (Figures 1C, S1E and S1F). Notochord formation is compromised in *tbxta* loss-of-function mutant embryos resulting in a dorsalization phenotype (no-tail) (Halpern et al., 1993; Schulte-Merker et al., 1994). Co-injection of RfxCas13d mRNAs with different sets of *tbxta* gRNAs (Figure 1C) recapitulated the no-tail phenotype at 30 hours post injection (Figures 1D, 1E and S1G). Importantly, the no-tail phenotype was only observed when *tbxta* gRNAs and *RfxCas13d* mRNA were co-injected (i.e., no phenotype was observed when either gRNAs or *RfxCas13d* mRNA were injected singly) and its penetrance correlated with *RfxCas13d* dosage (Figure 1E). Further, co-injection of *RfxCas13d* mRNA with gRNAs targeting *dnd1*, a gene controlling an unrelated developmental process (germ cell development and survival) (Weidinger et al., 2003), did not lead to dorsalization (Figure 1F). In addition, the CRISPR-RfxCas13d system triggered *tbxta* mRNA degradation by at least 6 hours post injection (Figure 1G). While the addition of nuclear localization sites (NLS) increased RfxCas13 activity in mammalian cells(Konermann et al., 2018), we observed that incorporation of an NLS decreased phenotypic penetrance in zebrafish embryos (Figure S1H) (P < 3.4e-9, Chi-Square). Finally, to confirm the specificity of the phenotype, we designed a new set of three gRNAs targeting the *tbxta* 3’UTR which also induced the no-tail phenotype (Figure 1F). This phenotype was specifically rescued (P < 2.07e-3, Chi-Square) by injection of the cognate mRNA (lacking endogenous 3’UTR), but not by injection of mRNA encoding GFP (P < 0.02, Chi-Square) or the antisense *tbxta* coding region (P < 0.29, Chi-Square) (Figures 1F and S1I). Together, these results establish CRISPR-RfxCas13d as an efficient tool to trigger endogenous mRNA knock-down during zebrafish embryogenesis.

**Figure 1.**
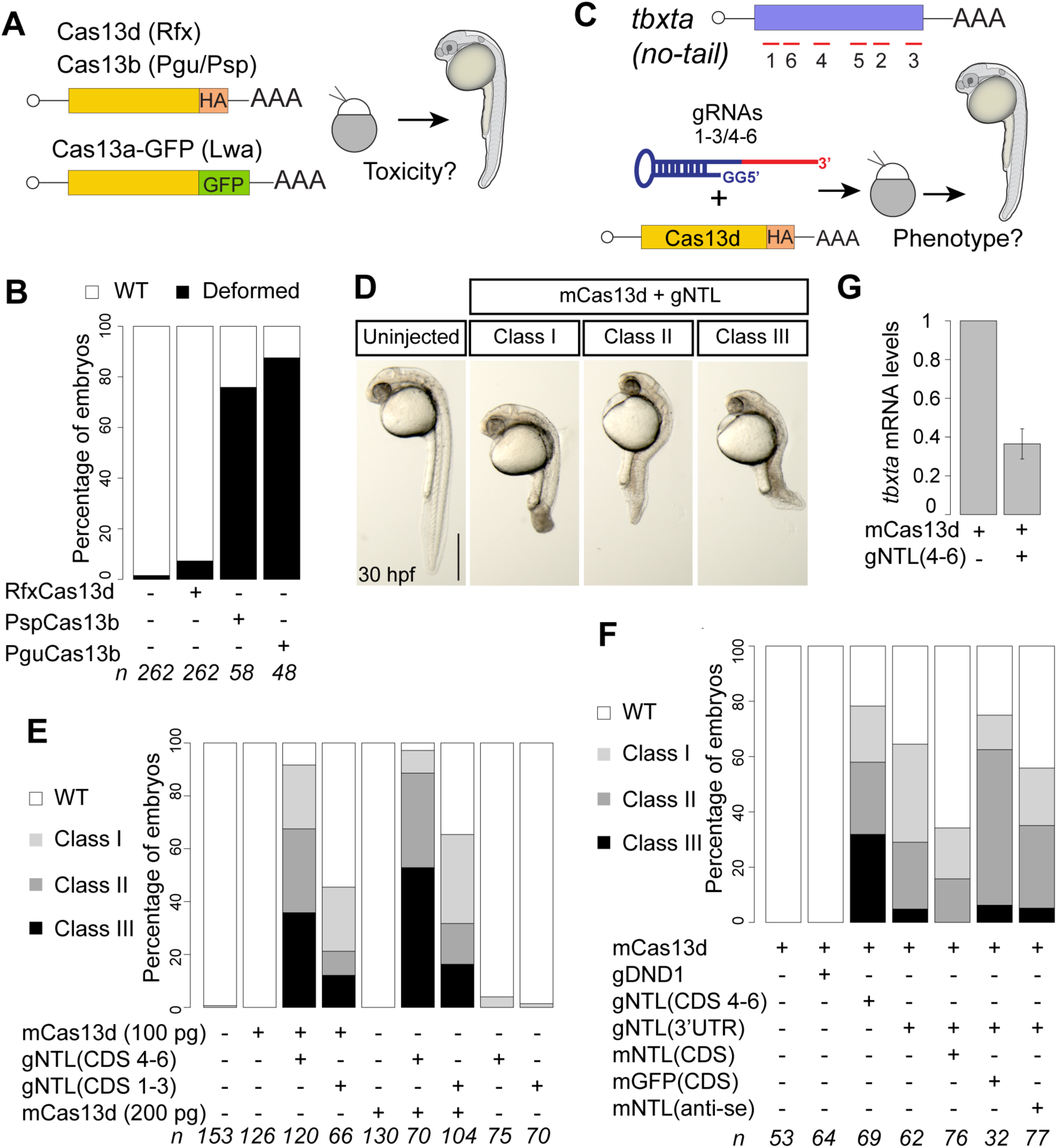
CRISPR-RfxCas13d system targeting *tbxta* recapitulates no-tail phenotype in zebrafish embryos. **A)** Schematic illustration of the experimental set-up used to analyze CRISPR-Cas13 toxicity. 200 pg of mRNA from each Cas13 variant was injected into one-cell stage zebrafish embryos. Cas13b and Cas13d are tagged with HA and Cas13a is fused to GFP. **B)** Toxicity evaluation of embryos. Stacked barplots showing the percentage of deformed and wild type (WT) zebrafish embryos (30 hours post fertilization, hpf) injected in conditions described in **a**. Number of embryos evaluated (n) is shown for each condition. **C)** Schematic of positions of six gRNAs (red lines) targeting *tbxta* CDS in zebrafish (top). Schematic of experimental set-up to analyze CRISPR-RfxCas13d-mediated RNA knockdown in zebrafish (bottom). Two sets of gRNAs targeting *tbxta* CDS (1-3 or 4-6; 300 pg per embryo) were mixed with mRNA (100 or 200 pg per embryo) coding for RfxCas13d and injected into one-cell stage embryos. **D)** Phenotypic severity in embryos injected with *RfxCas13d* mRNA (mCas13d) and gRNAs targeting *tbxta* (gNTL) compared to uninjected wild type (WT) evaluated at 30 hours post fertilization (hpf) (scale bar, 0.5 mm). Class I: Short tail (least extreme). Class II: Absence of notochord and short tail (medium level). Class III: Absence of notochord and extremely short tail (most extreme). **E)** Different gNTL sets recapitulate *no-tail* phenotype. Stacked barplots showing percentage of observed phenotypes under various injection conditions. Number of embryos evaluated (n) is shown for each condition. **F)** *no-tail* phenotype can be rescued. Stacked barplots showing percentage of observed phenotypes using gNTLs targeting 3’UTR (300 pg/ embryo) and mCas13d (200 pg/embryo) co-injected with different mRNAs: GFP (mGFP ORF, 100 pg/embryo), *tbxta* ORF with 3’UTR from pSP64T plasmid (mNTL CDS, 100 pg/embryo), or *tbxta* antisense (mNTL anti-se, 100 pg/ embryo) ORF. As an additional control, an unrelated gRNA set targeting a gene involved in germ cell formation (*dnd1*) was included (300 pg/embryo). Number of embryos evaluated (n) is shown for each condition. **G)** qRT– PCR analysis showing levels of *tbxta* mRNA at 7 hours post injection (hpi) in conditions described in panel **c**. Results are shown as the averages ± standard error of the mean from 2 independent experiments with 1 (mCas13d alone) or 2 biological replicates per experiment (n=20 embryos/ biological replicate) or Cas13 and Cas13 plus gNTL, respectively). p value of 0.0092 (t-test). *cdk2ap* mRNA was used as normalization control.

### CRISPR-RfxCas13d triggers effective and precise reporter mRNA depletion in zebrafish embryos

To further assess the efficacy of the CRISPR-RfxCas13d system to impair mRNA activity, two sets of three gRNAs each, specific to the CDS of either Red Fluorescent Protein (RFP) or Nano-luciferase mRNAs were individually injected along with *RfxCas13d* mRNA into zebrafish embryos (Figures 2A and B). Reporter activity (Figures 2C and D), as well as the respective mRNA levels (Figures 2 E and F), were significantly reduced in zebrafish embryos after 6 hours post injection when compared to embryos injected with control conditions (Figures 2C-F). Moreover, *RfxCas13d* mRNA coupled with individual gRNAs was sufficient to reduce RFP fluorescence intensity (Figure S2A). Notably, fluorescence intensity and activity of two control mRNAs, Green Fluorescent Protein (GFP) and Firefly, respectively, did not change appreciably in all conditions tested (Figures 2C, S2A and S2B), and only targeted mRNAs (RFP and Nano-luciferase) displayed reduced mRNA levels (Figures 2C-F and S2A), suggesting that the CRISPR-RfxCas13d system is specific.

**Figure 2.**
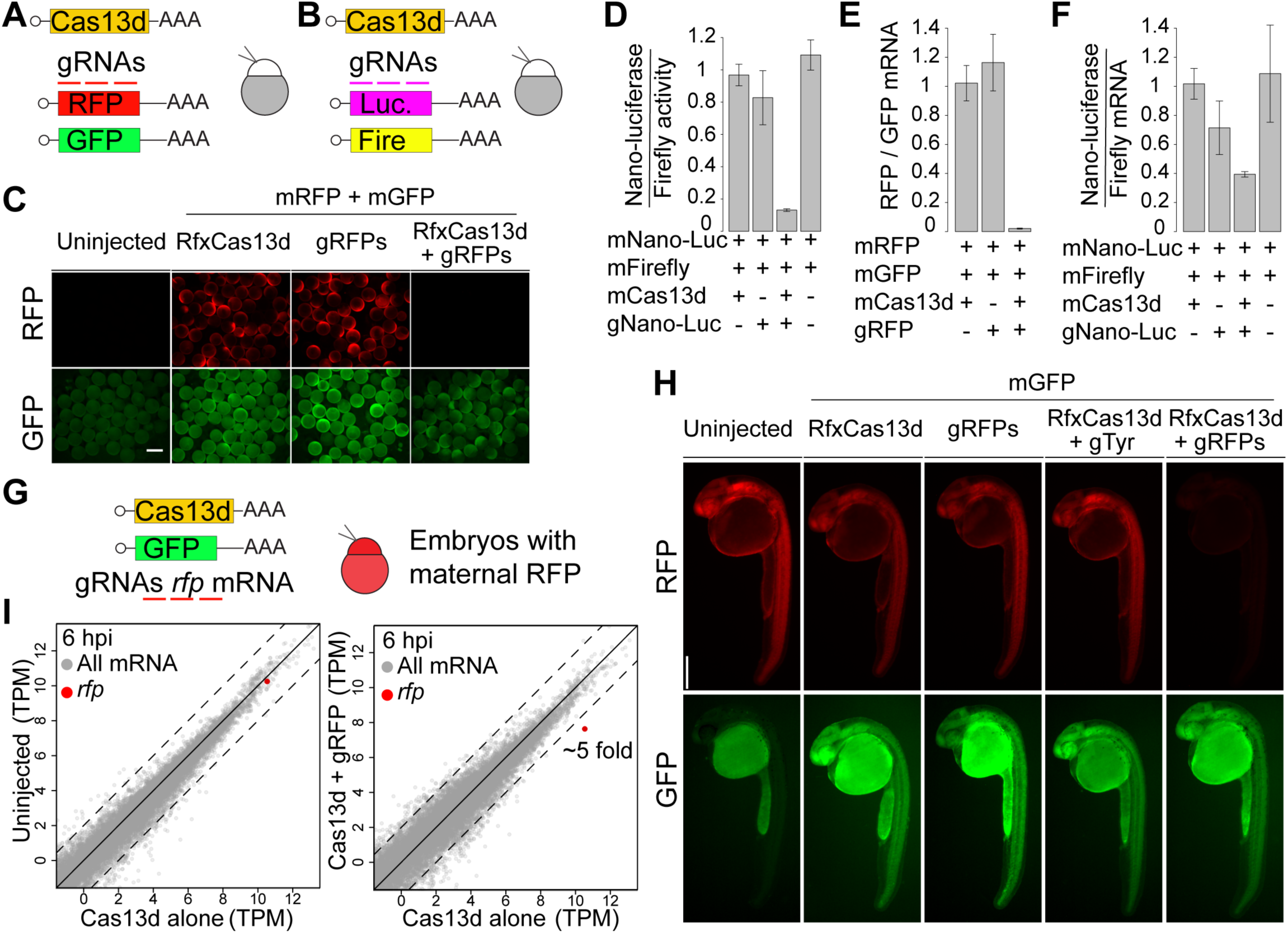
CRISPR-RfxCas13d specifically targets reporter mRNAs in zebrafish embryos. Schematics of **A**) *rfp* and *gfp* mRNAs or **B**) Nano-luciferase and Firefly mRNAs (10 pg/embryo) co-injected with/without *RfxCas13d* mRNA (200 pg/embryo) and/or gRNAs (250-400 pg/embryo) targeting *rfp* or *Nano-Luciferase* mRNA, respectively. **C)** Fluorescence images of representative embryos at 6 hpi with indicated mRNAs and gRNAs (scale bar, 0.5 mm). **D)** Ratio of nanoluciferase activity normalized to firefly luciferase activities under indicated conditions at 6 hpi. Results are shown as the averages ± standard error of the mean from at least 4 biological replicates per experiment (n=5 embryos/ biological replicate). p value < 0.0001, mCas13 and gNano-Luc compared to other conditions (one-way ANOVA). **E)** qRT-PCR analysis of ratio of RFP reporter mRNAs under indicated conditions at 6 hpi normalized with GFP mRNA. Results are shown as the averages ± standard error of the mean from 2 independent experiments with at least 2 biological replicates per experiment (n=20 embryos/ biological replicate). p value ≤ 0.0001, mCas13 and gRFP compared to other conditions (one-way ANOVA). **F**) Nanoluciferase mRNA level normalized to firefly mRNA level at 6 hpi at indicated conditions. Results are shown as the averages ± standard error of the mean from 2 independent experiments with 2 biological replicates per experiment (n=20 embryos/ biological replicate). p value < 0.05, mCas13 and gNano-Luc compared to mCas13d alone or mNano-Luc and mFirefly alone (one-way ANOVA) **G)** Schematic of mRNAs (*RfxCas13d* and *gfp*) and gRNAs targeting *rfp* introduced into embryos derived from transgenic mother expressing *rfp*. Fluorescence images of representative embryos at 26 hpi with indicated mRNAs and/or gRNAs. **H)** Fluorescence images of representative embryos at 26 hpi injected with indicated mRNAs and/or gRNAs (scale bar, 0.5 mm). **I)** Scatter plot showing mRNA level (RNA-seq) of zebrafish embryos at 6 hpi. Left panel, mRNA level of uninjected embryos versus embryos injected with *RfxCas13d* mRNA alone. Right panel, mRNA level of embryos injected with *RfxCas13d* mRNA plus rfp gRNAs versus embryos injected with *RfxCas13d* mRNA alone. *rfp* mRNA is indicated in red. Dashed lines indicate a 4-fold difference between RNA levels.

To address whether the CRISPR-RfxCas13d system could be used to target maternally deposited mRNAs, zebrafish embryos derived from a transgenic mother that expresses *rfp* at very high levels (i.e. among the top 100 most highly expressed maternal mRNAs) were injected with *RfxCas13d* mRNA plus three gRNAs targeting *rfp* (Figure 2G). Fluorescence intensity was visibly reduced after 24 hours post injection when compared to embryos injected with *RfxCas13d* mRNA or gRNAs alone, or by co-injection of *RfxCas13d* with unrelated gRNAs (Figure 2H). Interestingly, the reduced level of RFP persisted for the first 3.5 days and partially recovered after 5 days (Figure S2C and S2D, respectively). Furthermore, *rfp* mRNA levels displayed an approximately 4.4. fold reduction (77.3 %; P adjusted < 7.64e-6) compared to embryos injected with *RfxCas13d* mRNA alone after 6 hours post injection by RNA-seq (Figure 2I) and q-RT-PCR (Figure S2E). Moreover, RNA-seq analysis displayed a similar expression RNA level in zebrafish embryos injected with *RfxCas13d* mRNA alone or uninjected embryos (Figures 2I and S2F). Interestingly, within the co-injected embryos with *RfxCas13d* mRNA and gRNAs targeting *rfp* mRNA, RFP was the most down-regulated at the mRNA level (Figures 2I and S2F). Together, these results indicate that the CRISPR-RfxCas13d system can functionally disrupt the activity of maternally-provided mRNAs in a specific manner during early development in zebrafish embryos with minimal off-target effects detected.

### CRISPR-RfxCas13d system efficiently depletes endogenous maternally provided and zygotically-expressed mRNAs in zebrafish embryos

We next tested our optimized CRISPR/RfxCas13d method on endogenous mRNAs with known roles in embryonic development. We chose three genes whose loss-of-function phenotypes had been previously established. First, we targeted *dnd1* mRNA, whose impairment abolishes primordial germ cell migration and survival (Weidinger et al., 2003). Embryos co-injected with *RfxCas13d* mRNA and 3 gRNAs targeting *dnd1* were devoid of detectable germ cells, as indicated by loss of germ cell-specific GFP expression (from co-injection with GFP-nanos 3’UTR marking germ cells) (Figure 3A), recapitulating the phenotype observed in embryos injected with a morpholino targeting *dnd1* mRNA (Ciruna et al., 2002). In contrast, germ cells were not visibly affected by injection of *RfxCas13d* mRNA alone or in conjunction with gRNAs targeting an unrelated mRNA (Figure 3A). RNA-seq and qPCR analysis confirmed specific reduction (8-fold change −87.5%-, P adjusted < 2.76e-6, RNAseq) of *dnd1* mRNA levels (Figures 3B and S3A). Notably, *dnd1* mRNA was the most differentially expressed mRNA (at 6 hpi) when compared to embryos injected with *RfxCas13d* mRNA alone (Figures 3B and S3B). Together, these results demonstrate the robustness and specificity of *dnd1* knock-down by CRISPR-*RfxCas13d* system, which recapitulates the loss of germ cells.

**Figure 3.**
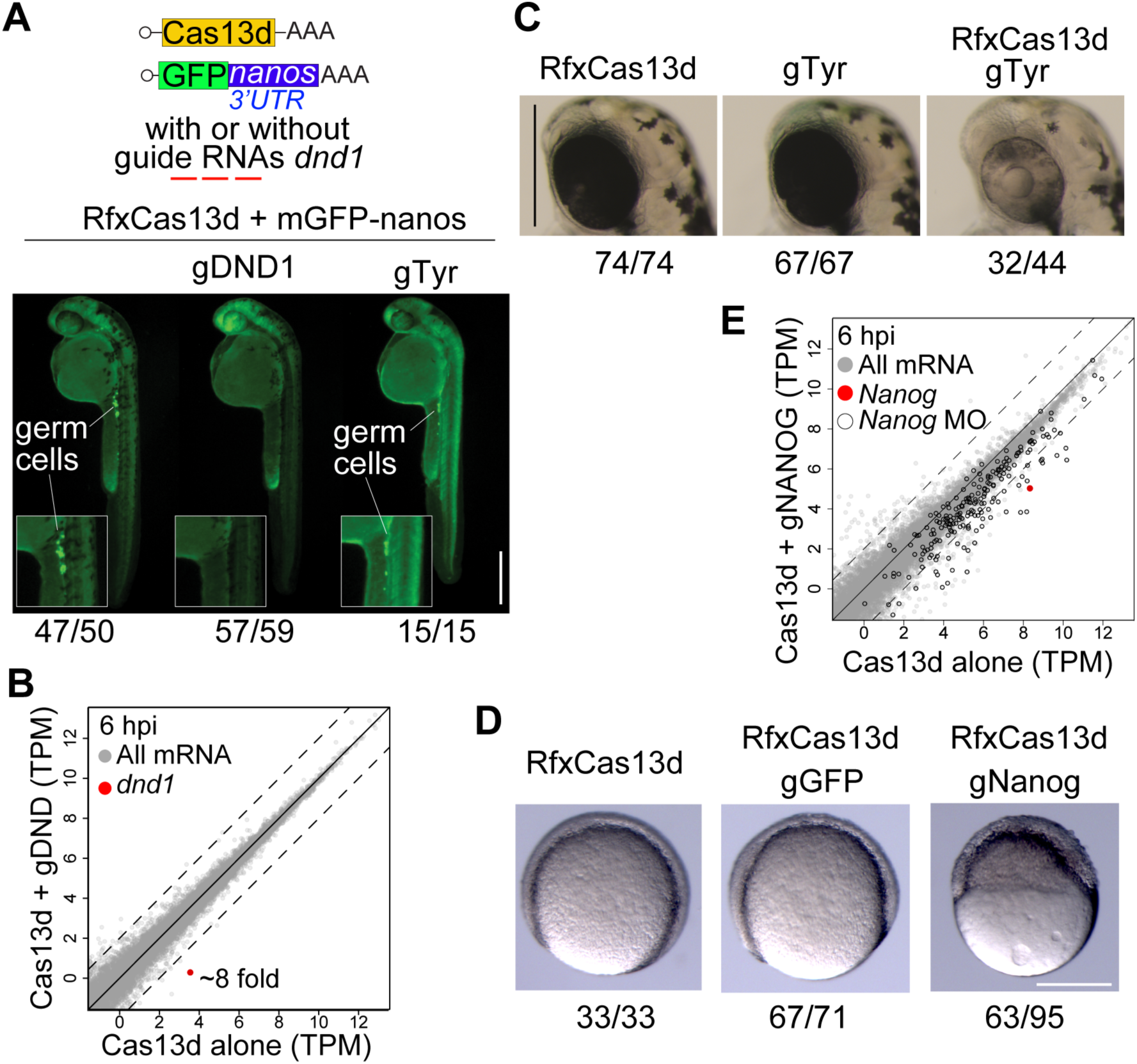
CRISPR-RfxCas13d recapitulates known loss-of-function zebrafish phenotypes. **A)** Schematic of *RfxCas13d* mRNA (200-250 pg/embryo) and *gfp* mRNA containing the 3’UTR of *nanos* (40 pg/embryo) co-injected with/without gRNAs targeting *dnd1* (500-600 pg/embryo) or *tyr (*500-600 pg/embryo). Germ cells are highlighted by germ-cell specific fluorescence (due to *nanos*-3’UTR) at 30 hpf. The ratio of embryos displaying intact germ cells (mCas13d alone or mCas13d plus gTyr) or without germ cells (mCas13d plus gDND1) vs total number of analyzed embryos (at least from two independent experiments) are indicated (scale bar, 0.5 mm). **B)** Scatter plot showing mRNA level (RNA-seq) of zebrafish embryos injected with *RfxCas13d* mRNA (250 pg/embryo) alone versus *RfxCas13d* mRNA plus *dnd1* gRNAs (600 pg/embryo) after 6 hpi, *dnd1* mRNA is indicated in red (~8-fold change). Dashed lines indicate a 4-fold difference between RNA levels. **C)** Pigmentation phenotypes (52 hpf) of head/eyes of embryos injected with *RfxCas13d* mRNA alone (300 pg/embryo, left), *tyr* gRNAs alone (600 pg/embryo, center), or both (right) (scale bars, 0.5 mm). The ratios of embryos displaying intact pigmentation (mCas13d or gTyr alone) or with the lack of pigmentation (mCas13d plus gTyr) vs total number of analyzed embryos are indicated. **D)** Representative pictures of epiboly-stage zebrafish embryos displaying gastrulation and epiboly defects in embryos injected with *RfxCas13d* mRNA plus *nanog* gRNAs (gNanog), but not in embryos injected with either *RfxCas13d* mRNA alone or *RfxCas13d* mRNA plus *gfp* gRNAs (gGFP). The ratios of embryos displaying normal development (*RfxCas13d* mRNA alone or *RfxCas13d* mRNA plus gGFP) or with epiboly and gastrulation defects (*RfxCas13d* mRNA plus gNanog) vs total number of analyzed embryos are indicated. Embryos were injected with 250 pg, 430 pg and 600 pg of mCas13d, gGFP and gNanog, respectively (scale bar, 0.1 mm). **E)** Scatter plot showing mRNA level (RNA-seq) of zebrafish embryos injected with *RfxCas13d* mRNA plus *nanog* gRNAs compare to embryos injected with *RfxCas13d* mRNA alone at 6 hpi. *nanog* mRNA is indicated in red. Black outline indicates genes that display reduced expression (>4-fold decrease) in *nanog* morhpholino-treated (MO) embryos. Dashed lines indicate a 4-fold difference between RNA levels.

Second, to test whether the CRISPR-RfxCas13d system could be used to reduce mRNA levels of later developmental genes, we designed three gRNAs targeting the tyrosinase (*tyr*) mRNA involved in pigmentation whose transcription starts at 16-17 hours post fertilization (Camp and Lardelli, 2001). Embryos injected with *tyr* gRNAs and *RfxCas13d* mRNA displayed loss of both eye and body pigmentation when compared to embryos injected with either *Rfx-Cas13d* mRNA or *tyr* gRNAs alone (Figures 3C and S3C). These results suggest that CRISPR-RfxCas13d can be used to knock-down zygotically-derived mRNAs transcribed later during embryogenesis.

Third, *nanog*, is a potent transcription factor involved in the activation of the zygotic gene expression in zebrafish (Gagnon et al., 2018; Lee et al., 2013; Veil et al., 2018) and injection of morpholinos targeting *nanog* mRNA or maternal and zygotic loss-of-function mutants induce severe gastrulation deficiencies and epiboly failure in zebrafish embryos (Lee et al., 2013; Veil et al., 2018). In a similar manner, embryos injected with *RfxCas13d* mRNA and three gRNAs targeting *nanog* mRNA displayed a significant reduction in *nanog* mRNA levels at 6 hours post injection (6-fold change, 84%, P adjusted < 3.536e-9, RNAseq) (Figures 3E, S3F and S3G) with similar consequences to gastrulation (Figures 3D, S3D and S3E) and downstream gene expression (*nanog* targets; Figure 3E). For example, mRNAs downregulated in embryos injected with *RfxCas13d* mRNA plus *nanog* gRNAs (139 mRNAs, P < 1e04, fold change> 2) were also downregulated in *nanog* morpholino-treated embryos (Lee et al., 2013) (P < 4.4e-11, Wilcoxon rank sum test) (Figures S3G and S3H). Conversely, mRNAs that displayed a greater than 4-fold reduction in *nanog* MO-treated embryos were also repressed after co-injection of *RfxCas13d* mRNA and *nanog* gRNAs (P < 1.7e-05, Wilcoxon rank sum test) (Figure 3E). Given the strong correspondence between MO and CRISPR-RfxCas13d methods, affected mRNAs are likely true targets of *nanog* (both direct and indirect). Altogether, our data show that the CRISPR-RfxCas13d system is a precise tool to impair maternal and zygotic gene activity in zebrafish embryos reducing in approximately 80% the level of mRNA (Figure S3I).

### CRISPR-RfxCas13d as RNA targeting tool in different animal models

To determine how the CRISPR-RfxCas13d system performs in other animal models, we first assessed its activity in medaka, another fish model for which RNAi is ineffective (Chen et al., 2017). Loss-of-function mutations in the *rx3* (*rax*) gene result in arrested eye development (Loosli et al., 2001). Co-injection of *RfxCas13d* mRNA plus three gRNAs targeting the *rx3* mRNA into one-cell stage medaka embryos resulted in *rx3* mRNA depletion (Figure S4A) (uninjected or *RfxCas13d* mRNA alone vs *RfxCas13d* mRNA and gRX3, P value=0.01 and 0.01, respectively; One-way ANOVA), and severe microphtalmia and/or anophthalmia, which were not observed with injection of *RfxCas13d* mRNA alone (Figures 4A, S4B and C) (P < 2.7e-12, Chi-Square). Next, we assessed the ability of RfxCas13d to disrupt reporter mRNA activity in the killifish (*Nothobranchius furzeri*), a research organism used to study aging and regeneration. We introduced mRNAs encoding *gfp*, *rfp*, *RfxCas13d*, as well as three gRNAs targeting *rfp*. We observed between 70 to 95% reduction in RFP mRNA level and fluorescence intensity 14 hours post injection when compared to control embryos (i.e. *RfxCas13d mRNA* or gRNAs alone; Figures 4B, S4D and S4E). Together, these results suggest that the CRISPR-RfxCas13d system can be widely applied across several animal species, including those historically recalcitrant to RNAi-based knock-down methodologies (e.g. zebrafish, medaka and killifish) (Chen et al., 2017).

**Figure 4.**
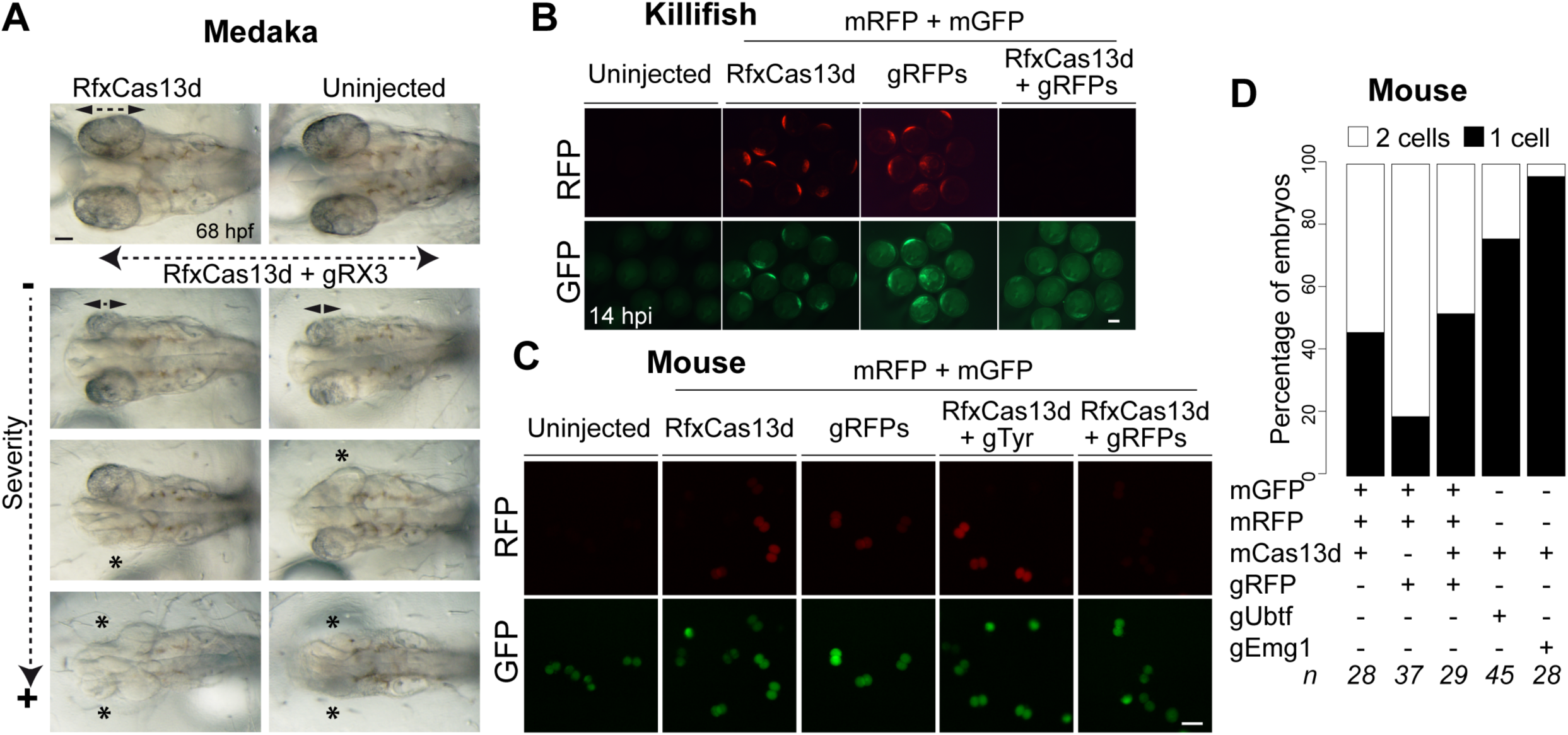
CRISPR-RfxCas13d as RNA targeting tool in different animal models. **A)** Representative pictures of medaka embryos at 68 hpf showing arrested eye development in embryos injected with *RfxCas13d* mRNA (450 pg) with gRNAs (1 ng) targeting *rx3* mRNA (gRX3) compared to the embryos injected with *RfxCas13d* mRNA alone or uninjected embryos. Double arrows and asterisk indicate eye length and absence of eye, respectively (scale bar, 0.1mm). **B)** Fluorescence microscopy images of killifish embryos showing RFP and GFP intensity of uninjected embryos or after 14 hpi injected with *rfp* and *gfp* mRNAs, with or without *RfxCas13d* mRNA (250-350 pg/embryo) and/or gRNAs (300 – 400 pg/embryo) targeting *rfp* mRNA (scale bar, 0.1 mm). **C)** Fluorescence microscopy images of mouse embryos 24 hours post injection (2 cell-stage) with *rfp* and *gfp* mRNAs with or without *RfxCas13d* mRNA and/or gRNAs targeting *rfp* or unrelated (zebrafish *tyr*) mRNA (scale bar, 100 um). **D)** Stacked plots showing the percentage of mouse embryos at 1 or 2 cells stage 24 hours post injection with the indicated mRNA and/or gRNAS. The number of embryos (n) is indicated from two or three independent experiments.

Although CRISPR-Cas13 has been shown to be effective in cultured mammalian cells, its efficacy has, to date, not been determined in a mammal *in vivo*. Therefore, we next assessed potential toxic effects of increasing concentrations of RfxCas13d mRNA in one-cell stage mouse embryos. We find that up to 1-2 pl of 25 ng/ul of *RfxCas13d* mRNA is tolerated with no deleterious effects (Figures S4F and S4G). Importantly, we observed lower RFP fluorescence intensity at 2-cell stage when *RfxCas13* mRNA was co-injected with *rfp* mRNA and respective gRNAs, compared to mouse embryos injected with either *RfxCas13* mRNA or *rfp* gRNAs alone, as well as in the presence of RfxCas13 mRNA plus three unrelated gRNAs targeting the *Danio rerio Tyr* gene (Figure 4C). To assess the effects on endogenous gene activity, we designed gRNAs targeting *ubtf* and *emg1* mRNAs. In both cases, introduction of *RfxCas13d* mRNA plus respective pools of gRNAs resulted in developmental arrest (P=1e-03 and P < 3.5e-05, respectively compared to *RfxCas13d*; Chi-Square) (Figure 4D), similar to previously observed phenotypes in loss-of-function mutant embryos (Hamdane et al., 2014; Wu et al., 2010). Together, these results suggest that the CRISPR-RfxCas13d system can serve as an effective additional knock-down tool in mice.

## Discussion

The ability to experimentally modulate gene activity is critical for understanding biological function. In this study, we show that the CRISPR-RfxCas13d system provides a robust, highly efficient, specific and straight-forward method to disrupt mRNA function in a wide variety of embryonic animal models.

A CRISPR-Cas-based system for perturbing RNA function has several advantages. Foremost, as shown here, Cas13-mediated knock-down can be applied to teleost fish models in which current RNAi methodologies remain ineffective (Chen et al., 2017; Kelly and Hurlstone, 2011). The CRISPR-Cas13 system does not burden endogenous cellular machinery. This is of particular concern in aquatic vertebrates such as *Xenopus*, where the sequestration of Argonaute proteins by exogenous RNAi is thought to compromise endogenous microRNA function (Kelly and Hurlstone, 2011; Lund et al., 2011). Another advantage lies in the ease, flexibility, and cost-effectiveness of this approach. Researchers can generate gRNAs through conventional *in vitro* transcription reactions with PCR-generated templates, allowing for both individual and pools of gRNAs. Since RfxCas13 does not have nucleotide sequence restrictions or protospacer flanking site (PFS) requirements in eukaryotic cells (Abudayyeh et al., 2017; Cox et al., 2017; Konermann et al., 2018), any sequence can be potentially targeted, including untranslated regions as well as long non-coding RNAs. Since mRNA levels are directly impacted by targeted degradation (in contrast to most MO-based strategies), validation can be easily performed via Northern blot analysis, quantitative RT-PCR, or RNA-seq. Moreover, the potential off-target effects can also be assessed by RNA-seq. In addition, we have found that RfxCas13d performs better than a RfxCas13d variant with two nuclear localization signals (NLS; on N- and C-termini) in zebrafish embryos. However, we cannot rule out the possibility that NLS-containing versions will be more effective in other model animal systems or under ubiquitous expression of both RfxCas13d and gRNAs, as has been shown in mammalian cells (Konermann et al., 2018).

Remarkably, our results indicate that injection of *RfxCas13d* mRNA plus gRNAs is sufficient to rapidly target maternally-provided *nanog* transcripts and recapitulates MO-induced molecular and developmental phenotypes, bypassing the need for using Cas13d purified protein. Given the ease and reduced cost of this approach, we envision the application of this tool toward large-scale screening of maternal factors (both coding and noncoding) involved in early development in different aquatic vertebrate systems. However, it is important to note that *in vitro* pre-assembly of the Cas13d protein complex may more rapidly reduce maternal mRNA activity and therefore, increase the penetrance of maternal phenotypes.

Nonetheless, we envision a range of potential future applications. Transgenic lines expressing RfxCas13d, either ubiquitously, or in an inducible, tissue or cell-specific manner, can be applied to the study of genes involved later in development, as well as for targeting genes whose disruption results in early lethality or deleterious effects of ambiguous origins. In sum, we have found that CRISPR-RfxCas13d system is adaptable and effective in a broad spectrum of animal models, and as such, will be an invaluable tool for unraveling gene function in a large variety of developmental contexts.

## Limitations

We have shown that CRISPR-RfxCas13d can disrupt maternal programs by targeting maternal mRNAs. However, as with other knock-down techniques in teleost embryos, this approach cannot target maternally provided protein. Further, although we find that injection of *rfxCas13d* mRNA is efficient in disrupting early maternal gene activity, (e.g.nanog), we posit that injection of purified RfxCas13d protein or maternal expression of RfxCas13d protein (via transgenic expression) could potentially lead to faster activation and thereby increase the penetrance of the maternal phenotype. Finally, it has recently been shown that genetic compensation can be triggered by RNA degradation (El-Brolosy et al., 2019; Ma et al., 2019). Further work should examine whether, in some cases, Cas13d-mediated cleavage of target mRNAs can activate the transcriptional adaptation response.

## Methods

### Guide RNA and mRNA generation

Guide RNA (gRNA) design was based on CRISPR-Cas13 targeting data in mammalian cell culture in which there was not any significant protospacer flanking site preference (PFS) reported and gRNA activity partially correlated with target accessibility (Abudayyeh et al., 2017; Konermann et al., 2018). Thus, mRNA targets were analysed *in silico* using RNAfold software (http://rna.tbi.univie.ac.at//cgi-bin/RNAWebSuite/RNAfold.cgi) (Lorenz et al., 2011) and protospacers of 22 nucleotides (target sequences) with high accessibility (low base-pairing probability from minimum free energy predictions) within the target mRNAs were selected to generate gRNAs. DNA template to generate gRNA was generated by fill-in PCR (Figures S1E and S1F). A gRNA universal primer (Table S1) containing the T7 promoter (5′-TAATACGACTCACTATA-3) and the processed direct repeat for RfxCas13d gRNAs (30 nt) preceded by 5′GG was used in combination with a specific oligo of 42 nt adding the spacer (22 nt for target-binding) and part of the repeat sequence (reverse complement orientation). A 71 (bp PCR product was generated following these conditions: 3 min at 95°C, 35 cycles of 30s at 95°C, 30s at 51°C, and 30s at 72°C, and a final step at 72 °C for 7 min. PCR products were purified using Favorprep Gel/PCR purification kit (Favorgen) columns or PCR purification kit (QIAgene) and used as template (300-500 ng) for a T7 in vitro transcription reaction (AmpliScribe-T7-Flash transcription kit from Epicenter; 12-16 h of reaction). *In vitro* transcribed gRNAs were DNAse-treated using TURBO-DNAse for 15 min at 37°C and precipitated with Sodium Acetate/Ethanol. gRNAs were visualized in a 2% agarose stained by ethidium bromide to check for RNA integrity and quantify using Qubit™ RNA BR Assay Kit (ThermoFisher, Q10210). gRNAs targeting *tbxta* and *tyr* CDS in zebrafish and *rx3* in medaka were *in vitro* transcribed from PCR containing a pool of different gRNA primers at equal concentration. All other gRNAs were individually *in vitro* transcribed and mix at equal concentrations in pools after the *in vitro* transcription. To increase mRNA depletion 3-4 gRNAs targeting the same mRNA were co-injected, otherwise is specified in figure legends.

Human codon-optimized RfxCas13d or NLS-RfxCas13d-NLS were PCR-amplified using primers Rfx13d_Fw and Rfx13d_Rv and Rfx13d-NLS_Fw and Rfx13d-NLS_Rv, respectively (Table S1) and using addgene plasmid # 109049 (a gift from Patrick Hsu) as template (Konermann et al., 2018). The following PCR products were then digested with *NcoI* and *SacII* and ligated into the pT3TS-nCas9n (Jao et al., 2013) plasmid previously digested with the same enzymes. Human codon-optimized PspCas13b and PguCas13b inserts were PCR-amplified from addgene plasmids #103862 and #103861 (a gift from Feng Zhang) (Cox et al., 2017) using primers PspCas13b_FW/RV (Table S1) and PguCas13b_FW/FR, respectively (Table S1). Both Cas13 PCR amplicons were subsequently digested using *EcoRI*-and *NotI,* then were cloned into vector pSP64T (Moreno-Mateos et al., 2017), which was previously digested with the same restriction enzymes. LwaCas13a-msfGFP, without and with NLS, was amplified from addgene plasmid #91902 (a gift from Feng Zhang) using primers Cas13aORF_Fw and Cas13aORF_Rv or Cas13aORF-NLS_Rv (Table S1), respectively. Both Cas13 PCR amplicons were subsequently digested using *EcoRI*-and *XhoI,* then were cloned into vector pSP64T. *tbxta* ORF (sense and antisense) was PCR amplified (Table S1). PCR products were digested with *EcoRI* and cloned into pSP64T previously digested with the same enzyme. p515-Tol2-attP-Ubi-RFP expression vector (see below RFP transgenic zebrafish section) was built by first cloning tag-RFP from p514-Ubi-TagRFP (a gift from Daniel Cifuentes of Boston University School of Medicine) into p328-attB-ubi:EGFP (gifted from the Piotrowski Lab at Stowers Institute) (Mosimann et al., 2011) via NEBuilder HiFi assembly kit (NEB) using PCR products from p328 plasmid amplified with pDest-gibson_FW/RV primers (Table S1) and from p514 plasmid amplified with tagRFP-Ubi_FW/RV (Table S1). Next the subsequent construct was digested using *KpnI* and *NotI*. Plasmid p511-Tol2-attP-Tol2, purchased from GENEWIZ (Kwan et al., 2007), was digested using the same enzymes and cloned into the p328 digest, resulting in the final expression plasmid used for transgenesis, p515-Tol2-attP-Ubi-RFP. All final constructs were confirmed by Sanger sequencing (Data S1).

To generate *tbxta*, *RfxCas13d* mRNA and *LwaCas13a-GFP* the DNA templates were linearized using *XbaI* and with *SmaI*, *SacI* to produce *PspCas13b*, *PguCas13b*, respectively. mRNA was synthesized using the mMachine T3 (*Cas13d*) or SP6 (*Cas13b*, *tbxta* and *Cas13a*) kit (Ambion). *In vitro* transcribed mRNAs were DNAse-treated using TURBO-DNAse for 15 min at 37°C and purified using the RNeasy Mini Kit (Qiagen) and quantifed using Qubit™ RNA BR Assay Kit (ThermoFisher, Q10210).

### Western blot

Ten embryos were collected at 6 hours post injection and transferred to 200 µl of deyolking buffer for washing (55 mM NaCl, 1.8 mM KCl, and 1.25 mM NaHCO3). Deyolking buffer was discarded and 200 µl of the same buffer were added to resuspend the embryos by pipetting. The resuspended embryos were incubated at room temperature for 5 min with orbital shaking, and then centrifuged at 300 g for 30 s and washed with 110 mM NaCl, 3.5 mM KCl, 10 mM Tris-HCl pH 7.4, and 2.7 mM CaCl. The pellet was resuspended in SDS-PAGE sample buffer before separation by SDS-PAGE and transferred to nitrocellulose membrane. Anti-HA antibody (11867423001, Roche) was used according to the manufacturer’s instructions. Secondary anti-mouse antibody (A5278, Sigma-Aldrich) was HRP-labelled and used according to the manufacturer’s instructions. Protein bands were visualised by triggering the HRP-catalysed chemiluminescent reaction catalysed through addition of the “Clarity” (TM) Western ECL Substrate (BioRad) following the manufacturer’s instructions, and recording the chemiluminescent images in a “ChemiDoc” (TM) MP Imaging System (BioRad). To verify the transfer of proteins to the membrane stain-free technology (Bio-Rad) was used.

### Zebrafish maintenance and image acquisition

All experiments involving zebrafish at CABD conform national and European Community standards for the use of animals in experimentation and were approved by the Ethical committees from the University Pablo de Olavide, CSIC and the Andalucian Government. Zebrafish wild type strains AB/Tübingen (AB/Tu) were maintained and bred under standard conditions (Westerfield, 1995). Wild-type zebrafish embryos were obtained through natural mating of AB/Tu zebrafish of mixed ages (5–18 months). Selection of mating pairs was random from a pool of 20 males and 20 females. Zebrafish embryos were staged in hours post-fertilization (hpf) as described (Kimmel et al., 1995). Zebrafish experiments at Stowers Institute were done according to the IACUC approved guidelines. Zebrafish embryos collected for microinjections were coming from random parents (AB, TF and TLF, 6-25 months old) mating from 4 independent strains of a colony of 500 fish. The embryos were pooled from random 24 males and 24 females for each set of experiment. Wild type fish lines from University of New Haven were maintained in accordance with OLAW and USDA research guidelines, under a protocol approved by the University of New Haven IACUC. Twenty couples of males and females (TU and AB intermixed strains, 6-18 month old) were randomly selected and crossed to generate embryos for experiments described herein. For each set of experiments from different laboratories both controlled and test embryos after injections were kept in similar condition and 200-800 embryos were injected per experiment (depending on the need of the experiment).

1-2 nl containing 100-300 pg of *cas13(s)* mRNA (Figure S1D) and 300-600 pg of gRNA were injected in the cell of one-cell stage embryos (see figure legends for details in each experiment). Phenotypes and samples were collected, analyzed and quantified between 6 hours and 5 days post injection depending on the experiment. Zebrafish embryo phenotypes and fluorescent pictures were analysed using an Olympus SZX16 stereoscope and photographed with a Nikon DS-F13 digital camera and images processed with NIS-Elements D 4.60.00 software or were imaged using Zeiss SteREO Lumar.V12 and Leica MZ APO stereomicroscopes.

### Transgenic RFP zebrafish

Transgenic zebrafish were kept according to the National Research Council’s and IACUC approved protocol 2016-1059 at the Stowers Institute for Medical Research zebrafish facility. Wild-type embryos obtained from breeding of adult AB-TU and TL-TLF strains were co-injected with 1 nl p515-Tol2-attP-Ubi-RFP (20 ng/ul) and capped transposase mRNA (50 ng/ul) (Kwan et al., 2007), at one-cell stage. Mosaic larvae demonstrating positive RFP fluorescence at 5 days post fertilization (dpf) were raised to 70 dpf and outcrossed to TL-TLF wild-type fish. These embryos were screened at 5 dpf for ubiquitous RFP fluorescence and positive individuals were raised to use for embryo production. F2 embryos coming from RFP positive mothers were injected with *RfxCas13d* mRNA and RFP gRNAs.

### RNA-seq libraries and analysis

mRNAseq libraries were generated from 500 ng of high-quality total RNA, as assessed using the Bioanalyzer (Agilent). Libraries were made according to the manufacturer’s directions for the TruSeq Stranded mRNA LP Sample Prep Kit (Illumina, Cat. No. 20020594) with TruSeq RNA Single Indexes Set A and B (Illumina, Cat. No. 20020492 and 20020493). Resulting short fragment libraries were checked for quality and quantity using the Bioanalyzer (Agilent) and Qubit Fluorometer (Life Technologies). Libraries were pooled, re-quantified and sequenced as 75bp single reads on a high-output flowcell using the Illumina NextSeq instrument. Following sequencing, Illumina Primary Analysis version RTA 2.4.11 and bcl2fastq2 v2.18 were run to demultiplex reads for all libraries and generate FASTQ files.

RNA-seq reads were aligned using STAR version 2.6.1c to *Danio rerio* reference genome danRer11 from University of California at Santa Cruz with RFP exogenmos sequence incorporated in its index. Transcripts were quantified using RSEM version 1.3.0 to calculate the transcript abundance ‘TPM’ (Transcript per Million). Fold change for each gene was calculated using edgeR version 3.24.3 after filtering genes with a count of 10 reads in at least one library. The resulting p-values were adjusted with Benjamini-Hochberg method using R function p.adjust

### Medaka maintenance, injection and image acquisition (pics of phenotypes)

All experiments involving medaka conform national and European Community standards for the use of animals in experimentation and were approved by the Ethical committees from the University Pablo de Olavide, CSIC and the Andalucian Government. The medaka wild type strain (iCab) was maintained and bred under standard conditions. 15-20 couples of males and females (4 months old) were randomly selected and crossed to generate embryos for experiments described herein. Embryos were injected at single cell stage also according to standard procedures (Masato Kinoshita et al., 2012). Embryos were staged in hours post-fertilization (hpf) as described (Iwamatsu, 2004). 2 nl containing 450 pg of *RfxCas13d mRNA* and 1 ng of gRNA were injected in one-cell stage embryos. Embryos phenotype was examined and photographed between 30 and 72 hpf using a stereoscope (SZX16–DP71, Olympus).

### Killifish Microinjection

Experiments at Stowers Institute were done according to the IACUC approved guidelines (2019-090). Random 3-to-4 months old individuals of killifish strain-inbred GRZ line, with opposite sex were crossed and embryos were pooled for each set of experiment. Three breeding tanks were used for embryos collection. Each tank contained one single male and 5 female. Approximately 200 killifish embryos were injected at single cell stage with ~2 nl containing 200 pg of *RfxCas13d* mRNA and 600 pg of gRNA. Embryos were held in 1 mm-wide trenches on a 1.5% agarose plate during injection. Embryos were kept in 1X Yamamoto’s embryo solution with 0.0001% methylene blue at 28° Celsius in the darkness and fluorescent pictures were imaged using Leica M205 FCA stereomicroscopes. GFP and RFP fluorescence from these images were quantified using Fiji (Image J) software.

#### Mouse embryo donors and injections

Immature C57BL/6J female mice (3-4 weeks of age) were utilized as embryo donors. The C57BL/6J females were super ovulated following standard procedures with 5 IU PMSG (Genway Biotech #GWB-2AE30A) followed 46 hours later with 5 IU hCG (Sigma #CG5) and subsequently mated to fertile C57BL/6J stud males (4-6 months of age). Females were checked for the presence of a copulatory plug the following morning as an indication of successful mating. Out of the 20 super-ovulated females, approximately 10 – 12 females were plugged and harvested for embryo microinjection for each set of experiment. One-cell fertilized embryos were collected from the oviducts of mated females at 0.5 days post coitus and placed in KSOM media in a CO2 incubator at 37° celsius, 5% CO2 until microinjection. CRISPR-RfxCas13d reagents were microinjected into both the pronucleus and cytoplasm (1-2 pl total) of one-cell embryos using previously described techniques (Nagi et al., 2003). In brief, microinjection was performed using a Nikon Eclipse Ti inverted microscope equipped with Eppendorf TansferMan micromanipulators, Eppendorf CellTram Air^→^ for holding of embryos, and Eppendorf FemtoJet^→^ auto-injector. A small drop of M2 media was placed on a siliconized depression slide and approximately 20-30 C57BL/6J oocytes were transferred to the slide for microinjection. The slide was placed on the stage of the microscope and oocytes were injected at 20x. Immediately following microinjection, the embryos were returned to the CO2 incubator in KSOM culture media and observed daily for embryo development. Animals were maintained in Lab Animal Service Facility (LASF) of Stowers Institute at 14:10 light cycle and provided with food and water *ad libitum*. Experimental protocols were approved by the Institutional Animal Care and Use Committee at Stowers Institute and were in compliance with the NIH Guide for Care and Use of Animals.

For toxicity testing, embryos were injected as described above with varying concentrations of *RfxCas13d* mRNA to test for toxicity. Injections concentrations were 2.5 ng/ul, 10 ng/ul, 25 ng/ul and 50ng/ul. A Stock solution of *RfxCas13d* mRNA was serially diluted using Tris Low EDTA microinjection buffer. Initial observations indicated the embryos responded normally to the microinjection. Embryos were cultured overnight as described above and observed the following morning for development. Embryos were returned to the incubator and observed each day to track the development to the 3.5 days post coitus blastocyst stage.

For *RfxCas13d* mRNA and gRNA Injections, reagents were prepared and delivered on ice at the desired injection concentrations. Injections concentrations were *RfxCas13d* mRNA (25 ng/ul), gRNA RFP (100 ng/ul), GFP and RFP mRNA (8-10 ng/ul). Fluorescent images were taken using Nikon ECLIPSE T*i* 2 microscope.

### qRT-PCR

For checking the level of targeted mRNA in zebrafish embryos, 20 embryos per biological replicate were collected for all zebrafish qRT PCRs at the described hours post injection in figures or legends and snap frozen in liquid nitrogen. Total RNA was isolated using standard TRIzol protocol as described in the manual (Life technologies). The cDNA was synthesized using SuperCript™ IV First-Strand Synthesis System, following the manufacturer’s protocol. To set up the real time qPCR for different genes, 1/25 cDNA dilution was used using forward and reverse primers per mRNA (10 µM; Table S1) in a 10 µl reaction. Samples were prepared as instructed for automated Freedom EVO® PCR workstation (Tecan) and run in QuantStudio 7 Flex Thermo cycler (Applied Biosystem)

Eight medak*a* embryos were collected at stage 16 (21 hpf) per biological replicate (except for WT uninjected that 5 embryos were collected). Total RNA was isolated from embryos injected using TRIzol reagent (Life Technologies). An amount of 500-1000 ng of purified total RNA was then subjected to reverse transcription using the iScript cDNA synthesis kit (Bio-Rad), following the manufacturer’s protocol. 4.5 microliters from a 1/25 dilution of the cDNA reaction was used to determine the levels of *rx3* mRNA in a 10 µl reaction containing 0.5 µl of forward and reverse primers (10 µM each; Table S1), using iTaq Universal SYBR Green Supermix (Bio-Rad) and a CFX connect instrument (Bio-Rad). PCR cycling profile consisted in a denaturing step at 95 °C for 30 s and 40 cycles at 95 °C for 1 s and 60 °C for 30 s. Two technical qPCR replicates were performed per biological replicate.

### Nano-luciferase activity

For nano-luciferase mRNA targeting, 10 pg of nano-luciferase mRNA and firefly luciferase mRNA (as an internal control) together with different experimental conditions including RfxCas13d (200 pg) and/or gRNA-nanoluciferase cocktail (400 pg) were injected per zebrafish embryo at one cell stage. Injected embryos were collected at 6 hours post injection and at least 8 tubes (5 embryos per tube) per condition were snap frozen in liquid nitrogen. Nano-Glo Dual-Luciferase® reporter assay protocol (Promega) was followed as instructed in the manual for assaying the activity of nanoluciferase and firefly luciferase activity using infinite M200 PRO by TECAN.

### Statistics

No statistical methods were used to predetermine sample size. The experiments were not randomized, and investigators were not blinded to allocation during experiments and outcome assessment. No data were excluded from the analysis. Unpaired two-tailed Mann–Whitney and χ2-test (chi-squared) tests were used to compare the results from CRISPR-RfxCas13d and Morpholino (MO) targeting *nanog* and different injections conditions, respectively. These tests were performed using R. For quantitative RT-PCR and luciferase activity p value was calculated using unpaired t-test or one-way ANOVA test using Prism (GraphPad Software, La Jolla, CA, USA) and Standard Error of the Mean (SEM) was used to show error bars. As it was described in the RNA-seq section, the fold change for each gene was calculated using edgeR version 3.24.3 after filtering genes with a count of 10 reads in at least one library. The resulting p-values were adjusted with Benjamini-Hochberg method using R function p value adjusted using two biological replicates.

### Data availability

Original data underlying this manuscript can be accessed from the Stowers Original Data Repository at http://www.stowers.org/research/publications/libpb-1454. The Stowers Institute contributing authors have selected to present the raw data of this manuscript resulting from work performed at The Stowers Institute. The RNA-seq data have been deposited in GEO under accession code GSE135884. All the mRNA level measure by RNA-seq are provided in Table S2.

## Acknowledgements

We thank Raktim Roy and Kai Zhang and Hui Qian for initial support, Fish facility, Molecular Biology facility and Hassan Huzaifa and Shiyuan Chen from the Computational core facilities from Stowers Institute and Ana Fernández-Miñan, Laura Tomas and Alejandro Díaz from the CABD aquatic vertebrate and proteomic platforms and Genetics section within the Department of Molecular Biology and Biochemical Engineer at University Pablo de Olavide for reagents and experimental support. We also thank Paul Trainor and Michael Durnin for their help with mouse experiments. We thank Ignacio Maeso (CABD) and Alfonso Fernandez-Alvarez (CABD) and all members of the Moreno-Mateos and the Bazzini laboratories for intellectual and technical support. This work was supported by Ramon y Cajal program (RyC-2017-23041) and grants PGC2018-097260-B-I00 and MDM-2016-0687 from Spanish Ministerio de Ciencia, Innovación y Universidades and the Springboard program from CABD (M.A.M-M) and the Stowers Institute for Medical Research (AAB).

J.R.M-M. is supported by BFU2017-86339-P and MDM-2016-0687 grants (Spanish Ministerio de Ciencia, Innovación y Universidades). E. M-T and J.A.-NP are supported by INNOVATE PERÚ grant 168-PNICP-PIAP-2015 and FONDECYT travel grant 043-2019. The CABD is an institution funded by Pablo de Olavide University, Consejo Superior de Investigaciones Científicas (CSIC) and Junta de Andalucía.

## Author contributions

M.A.M.-M. and A.A.B. conceived the project and designed the research. M.A.M.-M., G. K., A.A.B. and J.A.-NP. performed all zebrafish experiments with the contribution of M.D.V. and J.R.G. C.M.T. and E.O.B. carried out CRISPR-LwaCas13a experiments. J.R.M-M. M.A.M.-M and J.A.-NP. performed medaka experiments. W.W, T.J.C, A.M.M and G.K carried out killifish and mouse experiments. E.M-T. and A.S.A. provided reagents and materials. M.A.M.-M., A.A.B. and G.K. performed data analysis and M.A.M.-M., C.M.T. and A.A.B. wrote the manuscript with input from the other authors. All authors reviewed and approved the manuscript

## Competing interests statement

The authors declare no competing non-financial interests.

## Supplementary Information

**Supplementary figure 1.**
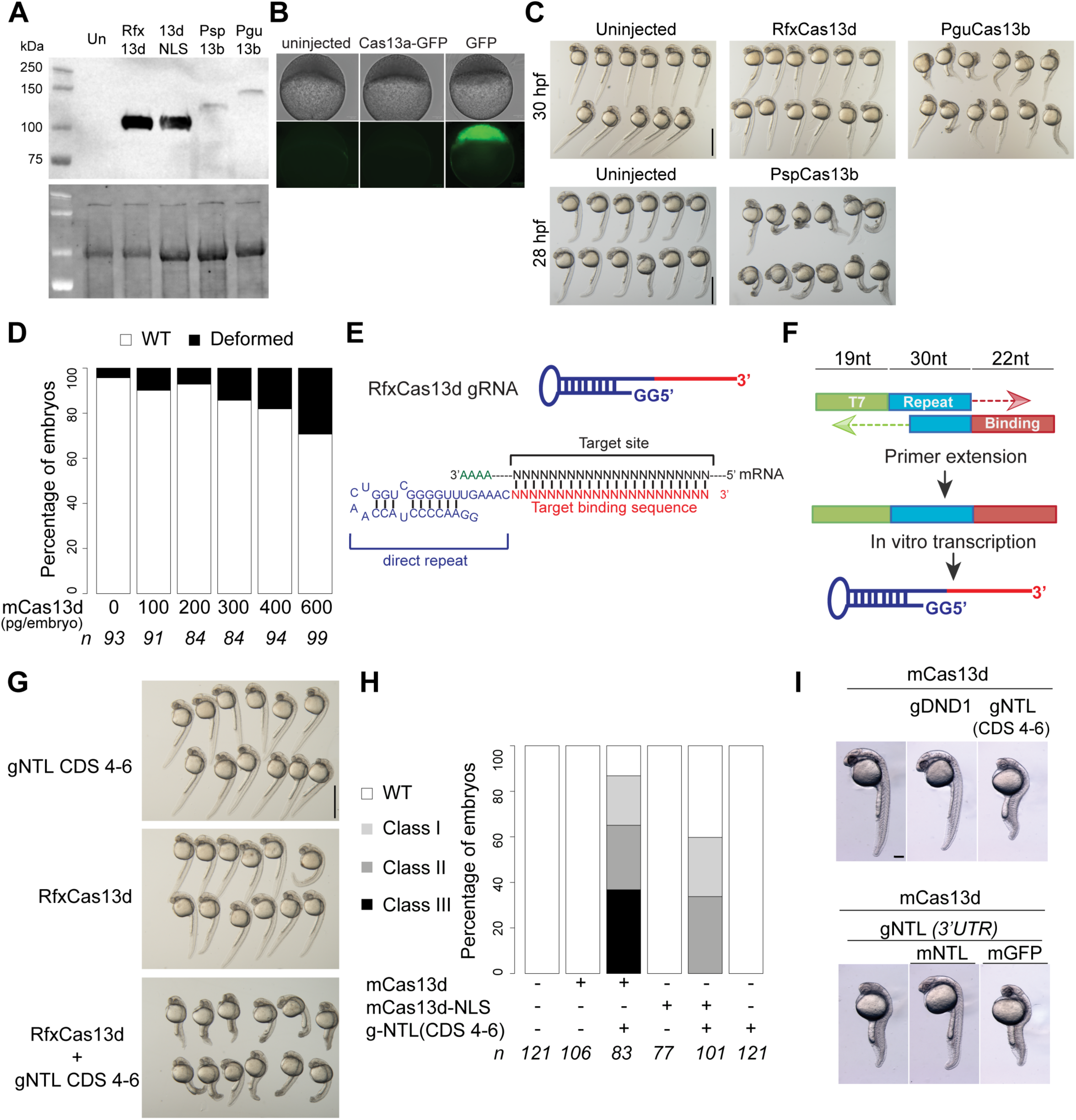
CRISPR-Cas13 optimization in zebrafish. **A)** Western-blot for HA showing protein level at 6 hpi of the indicated Cas13 after injection of the encoding mRNA for each respective *Cas13* mRNA into single cell zebrafish embryos. **B)** Representative pictures of embryos at 4.5 hours post fertilization (hpf) uninjected or injected with 200 pg of mRNA encoding for Lwa-Cas13a-GFP or GFP (scale bar, 0.1 mm). Lack of GFP expression was also observed using Cas13a-NLS-GFP version (not shown). **C)** Representative pictures of zebrafish embryos at 30 (top) or 26 (bottom) hpf uninjected or injected with 200 pg of mRNA encoding for RfxCas13d, PguCas13b or PspCas13b (scale bar, 1 mm).**D)** Toxicity evaluation of embryos. Stacked barplots showing the percentage of deformed and wild type (WT) zebrafish embryos (30 hpf) injected with different concentrations of *RfxCas13d* mRNA. Number of embryos evaluated (n) is shown for each condition. **E)** Schematic of gRNA structure for RfxCas13d (top) and scheme showing a gRNA (binding sequence in red, direct repeat in blue) binding to the mRNA (black). Adapted from Moreno-Mateos et al 2017^30^. **F)** Diagram of the gRNA generation by fill-in PCR followed by *in vitro* transcription. An oligonucleotide containing the T7 promoter (green) followed by two Guanine and 30 nt of the direct repeat (in blue) for annealing is used in combination with oligonucleotide containing 20 nt of the reverse complement of the repeat and 22 nt of the binding sequence (in red). Adapted from Moreno-Mateos et al 2017^34^. **G)** Representative pictures of zebrafish embryos at 32 hpf injected with gRNAs targeting *tbxta* (gNTL 4-6; 300 pg/embryo) and/or *RfxCas13d* mRNA (200 pg/embryo) (scale bar, 1 mm). **H)** Stacked barplots showing percentage of observed phenotypes using gNTLs targeting CDS (gNTLS 4-6, 300 pg/embryo) co-injected with *RfxCas13d* mRNA without (mCas13d) or with C- and N-terminal nuclear localization sites (mCAS13d-NLS) (200 pg/embryo each mRNA). Knockdown classes are described in Figure 1D. Number of embryos evaluated (n) is shown for each condition. **I)** Pictures of representative embryos injected with the indicated mRNA and/or gRNAs described in (scale bar, 0.1 mm) **Figure1F.**

**Supplementary figure 2.**
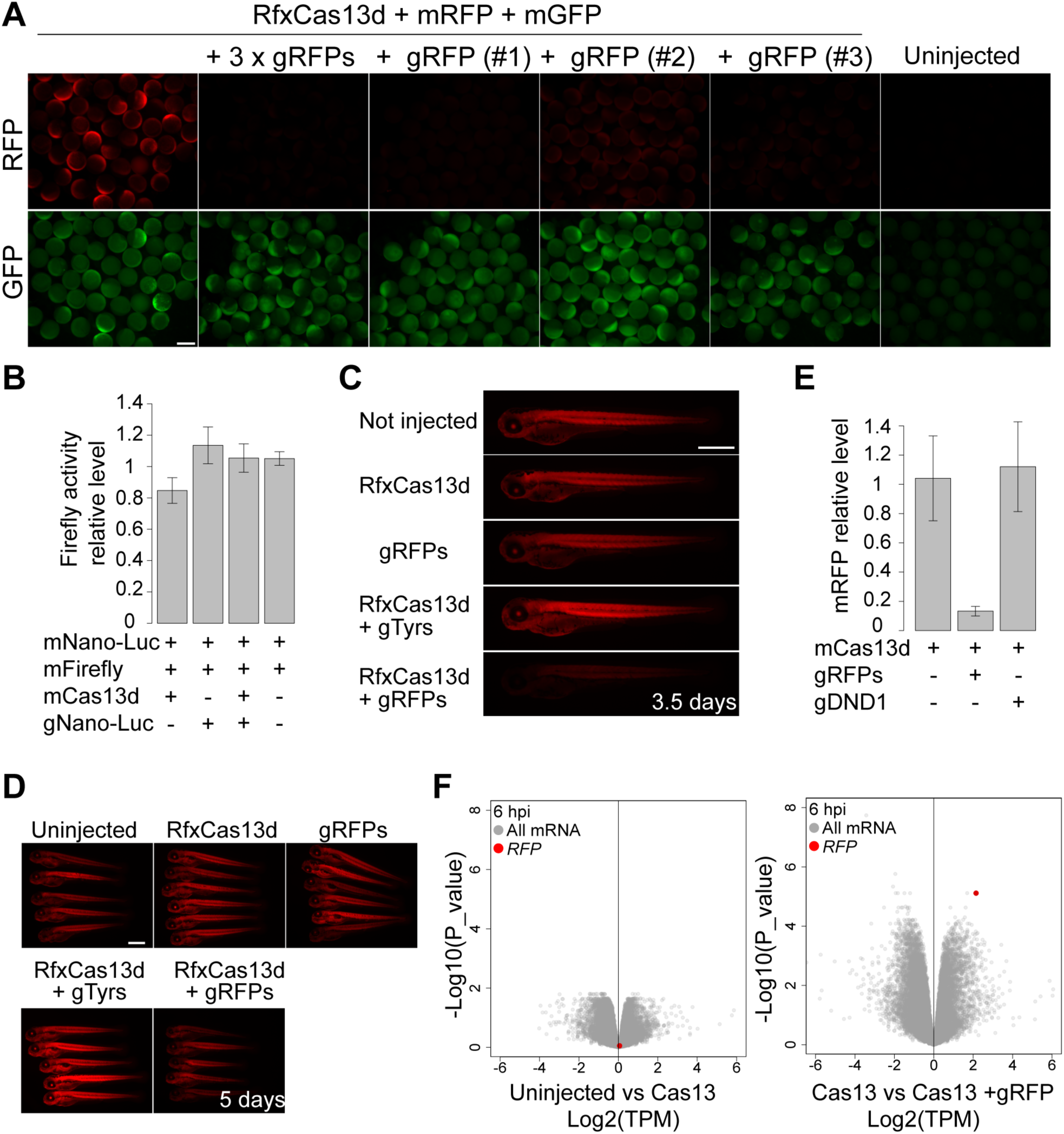
CRISPR-RfxCas13d system efficiently targets reporter mRNAs in zebrafish. **A)** Fluorescence microscopy images of zebrafish embryos at 6 hours post injection with the indicated mRNA and gRNAs (scale bar, 0.5 mm). **B)** Relative level of firefly luciferase activity at 6 hours from the firefly luciferase mRNA injected as an internal control for each combination described in Figure 2D**. C)** Fluorescence microscopy images showing the RFP intensity in gRNA targeted RFP in transgenic fish at 3.5 and **D)** 5 days post injection using the indicated mRNA and/or gRNAs, suggesting time dependent recovery of RFP expression (scale bar, 0.1 mm). **E)** qRT–PCR analysis showing levels of *rfp* mRNA at 6 hpi in transgenic embryos injected with *RfxCas13d* mRNA alone or together with gRNAs targeting *dnd1* or *rfp* mRNAs. Results are shown as the averages ± standard error of the mean from 2 independent experiments (n=20 embryos / experiment). *Taf15* mRNA was used as normalization control. **F)** Scatter plots representing the fold change in mRNA and the associated P value from two biological RNA-seq replicates at 6 hours post injection with the indicated gRNAs and/or mRNA encoding for RfxCas13d in RFP transgenic embryos. *rfp* mRNA is indicated in Red.

**Supplementary figure 3.**
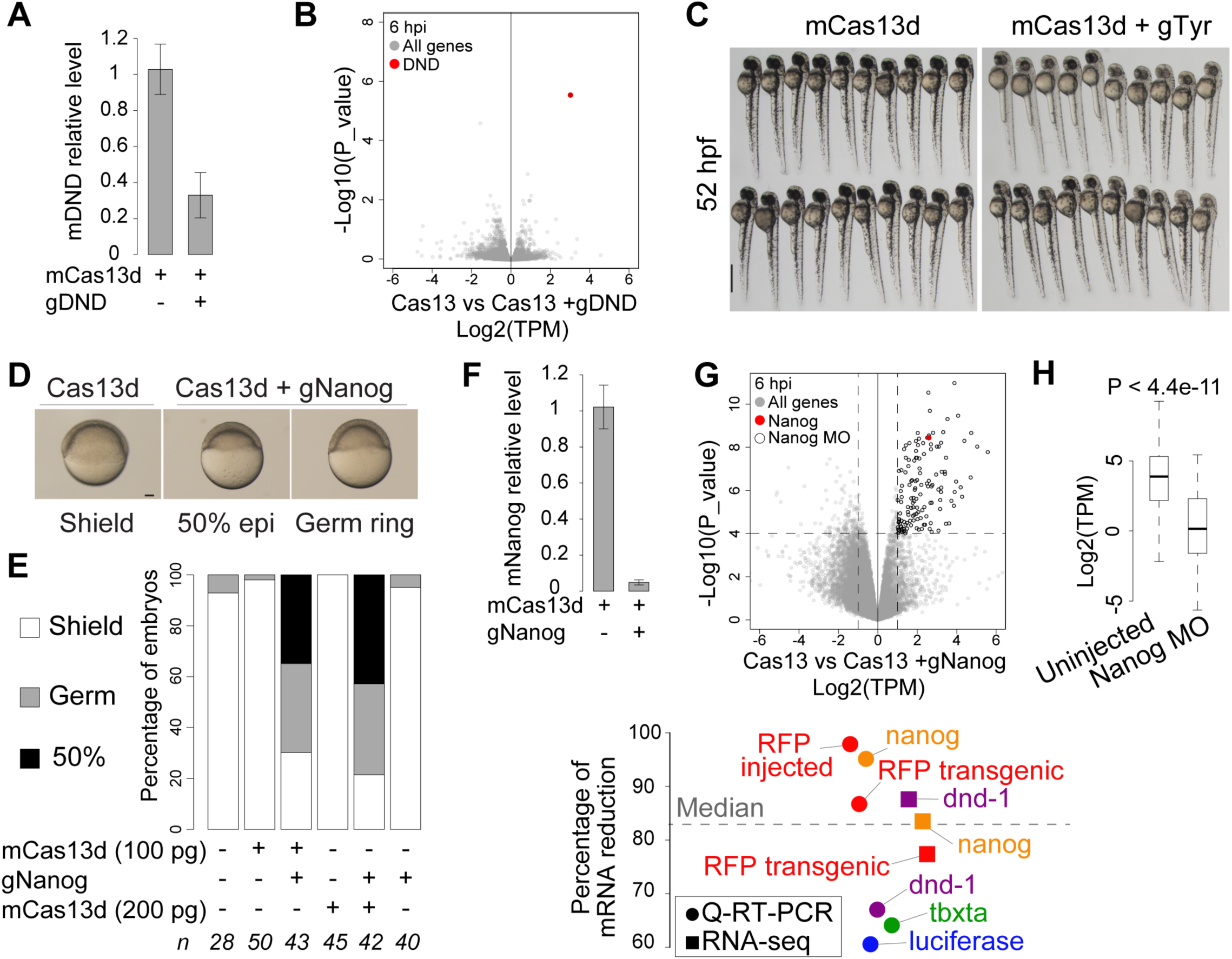
CRISPR-RfxCas13d system efficiently targets endogenous mRNAs in zebrafish. **A)** qRT–PCR analysis showing levels of *dnd1* mRNA in embryos at 6 hpi injected with *RfxCas13d* mRNA alone or together with gRNAs targeting *dnd1 mRNA.* Results are shown as the averages ± standard error of the mean from 2 independent experiments with 2 biological replicates per experiment (n=20 embryos/ biological replicate). *Taf15* mRNA was used as normalization control (P value=0.0099, t-test). **B)** Scatter plots representing the fold change in mRNA and the associated P value from two biological RNA-seq replicates at 6 hours post injection with gRNAs targeting *dnd1* and/or mRNA encoding for RfxCas13d. *dnd1* mRNA is indicated in Red. **C)** Representative pictures of zebrafish embryos from the experiment described in Figure 3C (scale bar, 1 mm). **D)** Representative pictures of embryos injected with gRNAs targeting *nanog* CDS (225 pg per embryo injected at one-cell stage, gNanog) and/or mRNA (100 or 200 pg per embryo injected at one-cell stage) coding for RfxCas13d (Cas13d) showing different developmental stages. Shield, germ ring and 50% epiboly (50% epi) correspond to 6, 5.7 and 5.3 hpf in wild type embryos growing in standard conditions, respectively (scale bar, 0.1 mm.) **E)** Stacked barplots showing percentage of observed phenotypes under conditions described in panel **e**. Number of embryos evaluated (n) is shown for each condition. **F)** qRT–PCR analysis showing levels of *nanog* mRNA in embryos at 6 hpi injected with *RfxCas13d* mRNA alone or together with gRNAs targeting *nanog* mRNA (P value < 0.0001, t-test). Results are shown as the averages ± standard error of the mean 2 independent experiments with at least 2 biological replicates each (n=20 embryos/ biological replicate). *Taf15* mRNA was used as normalization control. **G)** Scatter plots representing the fold change in mRNA and the associated P value from two biological RNA-seq replicates at 6 hours post injection with gRNAs targeting *nanog* and/or mRNA encoding for RfxCas13d. *nanog* mRNA is indicated in Red. Dashed lines indicate more than 4-fold differences between in RNA level and P adjusted value = 1e4). Black outline indicates mRNA that display reduced expression (>4-fold decrease and < P adjusted value = 1e4) due to the injection of RfxCas13d with gRNAs targeting *nanog*. **H)** Boxplot showing that the mRNAs down regulated in the embryos injected with RfxCas13d with gRNAs targeting *nanog* (Black outline in panel g) also display lower mRNA level in embryos injected with morpholino targeting *nanog* expression compare to uninjected embryos (P value < 4.4e- 11, Mann–Whitney *U*-test). The box defines the first and third quartiles, with the median indicated with a thick black line and vertical lines indicate the variability outside the upper and lower quartiles. **i)** The percentage of mRNA reduction due to CRISPR-RfxCas13d for each targeted mRNA measured by RNA-seq (square) or q-RT-PCR (circles). Dashed line indicates the median of all the mRNA knock-downs.

**Supplementary figure 4.**
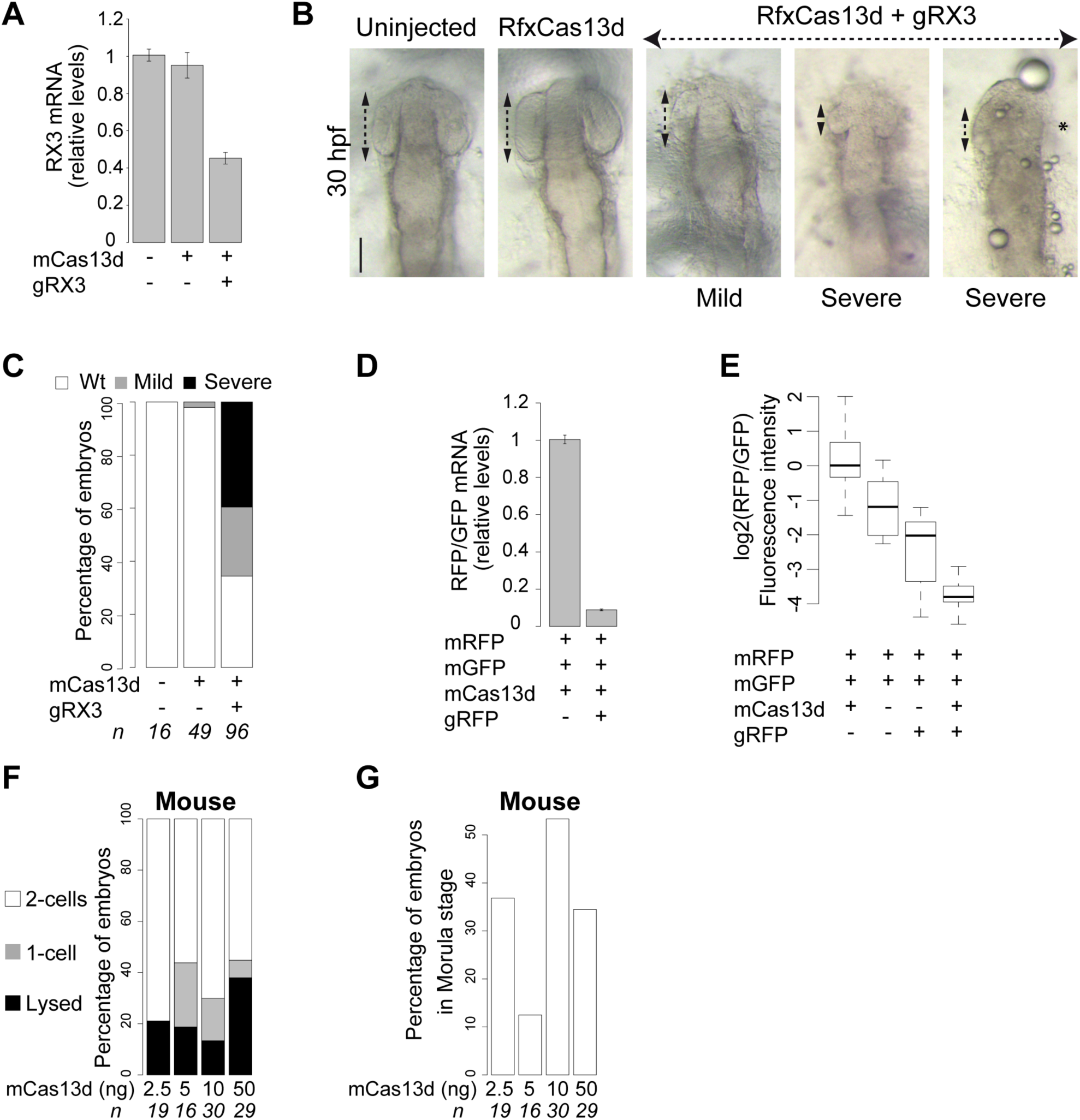
CRISPR-RfxCas13d targets different mRNAs in animal models. **A)** qRT–PCR analysis showing levels of *rx3* mRNA in medaka embryos at 21 hpf injected with *RfxCas13d* mRNA alone or together with gRNAs targeting *rx3* mRNA. Results are shown as the averages ± standard error of the mean from 2 (uninjected embryos) or 3 biological replicates (two technical replicates per biological replicate). *actin-beta* mRNA was used as normalization control. The data were subjected to one-way ANOVA test. **B)** Representative pictures of medaka embryos at 30 hpf injected in the conditions described in Figure 4A and showing different grade of severity of the phenotype. Double arrows and asterisk indicate eye length and absence of eye, respectively (scale bar, 0.1 mm). **C)** Stacked barplots showing percentage of observed medaka phenotypes at 30 hpf (**panel a**) under injection conditions described in Figure 4A. Number of embryos evaluated (n) is shown for each condition. **D)** qRT–PCR showing levels of *RFP* mRNA reduction with respect to mGFP in Killifish embryos injected with *RFP*, *GFP* and *RfxCas13d* mRNA with or without gRNAs targeting *RFP* mRNA. Results are shown as the averages ± standard error of the mean from 2 biological replicates (P= 0.0081, t-test) (n=20 embryos/biological replicate) **E)** Box plot showing the logarithmic ratio between the RFP and GFP fluorescence intensity observed in killifish embryos after 14 hpi with the indicated mRNA and/or gRNAs from at least 9 embryos per condition. The RFP/GFP intensity ratio was compared between mRFP-mGFP condition (without CRISPR-RfxCas13d) vs mCas13d alone, gRFP alone and mCas13d plus gRFP, (P< 0.0001, P=0.1366 and P= 0.0216, respectively; one-way ANOVA). The box defines the first and third quartiles, with the median indicated with a thick black line and vertical lines indicate the variability outside the upper and lower quartiles. **F)** Stacked plots showing the percentage of mouse embryos at 1 or 2 cells stage or lysed after 24 hours post injection with 1-2 ul of the indicated *RfxCas13d* mRNA concentration. The number of embryos (n) is indicated for each condition. **G)** Bar plot showing the proportion of embryos injected with different *RfxCas13d* mRNA concentration that reach Morula stage. The number of embryos (n) is indicated for each condition.

